# Evidence for common spike-based temporal coding of overt and covert speech in pars triangularis of human Broca’s area

**DOI:** 10.1101/2024.03.21.586130

**Authors:** Philémon Roussel, Florent Bocquelet, Stéphan Chabardès, Blaise Yvert

## Abstract

Broca’s area has long been described as a central region of cortical speech networks. Yet, its detailed role during speech production remains largely unknown and even sometimes debated. Recordings at the individual neuron level could help understand whether and how speech is encoded in this region but such data remain very scarce. Here we use direct intracortical recording in *pars triangularis* of human Broca’s area to show that the encoding of speech in this region relies not only on spike rates but also on the precise timing of action potentials within individual spike trains. First, we found that the overall spike rate of the whole population remained unchanged between periods of rest, overt and covert speech, but that individual firing rates of a few neurons fluctuated across these conditions. These fluctuations resulted in different overall population dynamics across conditions. Next, we also observed that the temporal arrangement of spikes within individual spike trains was not purely random but also signed which sentence was produced. By matching ensemble spike trains based on inter-spike intervals, it was possible to decode which sentence had been pronounced well above chance and with a comparable accuracy than when using spike counts. Moreover, the temporal code characterizing the overt production of individual sentences was found to be largely conserved when the same sentences were covertly imagined and enabled to decode cover sentences with an accuracy 75% higher than when considering spike counts. Altogether, these results suggest the existence of 2 modes of speech encoding in Broca’s area, one based on the modulation of individual firing rates and the other based on the precise temporal structure of individual spike trains, and that the latter type of encoding remains more largely conserved between overt and covert speech production.

## INTRODUCTION

Following the seminal report by Paul Broca about a patient suffering from aphemia after damage of the posterior part of his left inferior frontal region^1^, this part of the brain, hence commonly known as Broca’s area, has received tremendous attention in the field of speech and language research. This region is indeed considered as a key part of the brain language network^2–8^ that has been widely shown to be active during speech production^9–12^, speech planning^13^, speech processing^14–18^ and even non-speech tasks^19–21^. The two Brodmann areas BA44 (*pars operculari*s) and BA45 (*pars triangularis*) that compose Broca’s area have been shown to be differently involved in speech processing. While *pars opercularis* is consistently reported to be active in numerous speech tasks including simple reading and repetition of words^22^ or phonetic processing tasks^23^, *pars triangularis* tends to be more commonly associated with higher-level language processes such as syntactic^13,24^ or semantic^24,25^ computations and learning of correct selections of vocal responses^26^. This view is consistent with intracranial electrophysiological recordings reporting an encoding of semantic information more anterior in the inferior frontal gyrus than the encoding of phonological and articulatory information^27,28^, with the observation that damage to BA44 is a stronger predictor of speech impairment than damage to BA45^29^, and with direct intraoperative cortical stimulation studies reporting speech arrest upon stimulation of the premotor/*opercularis* area^30,31^, but not upon stimulation of *pars triangularis*^32,33^, which rather induces speech sequence deficits^30^. This high-level role of Broca’s area in language production may explain why speech deficits are only transiently observed after a cortical damage or a surgical resection of this region, and why in these cases language functions typically recover within a few months as long as the ventral sensory-motor and premotor cortices dedicated to the control of articulation^34^, as well as the anterior part of the arcuate fasciculus^29,35,36^, remain intact.

Altogether, the current knowledge of the functional role of Broca’s area thus indicates that, compared to more posterior premotor and motor areas being dedicated to the motor execution of articulation, this area can be viewed as a key region for speech unification (i.e. the assembly of speech elements into meaningful sentences)^37,38^. This is further supported by studies using cortical cooling showing that silencing the ventral precentral gyrus or premotor cortex impacts the quality of speech^39,40^, while cooling *pars opercularis* and *pars triangularis* of Broca’s area stretches the timing of speech^40^.

This body of evidence suggests that the neural dynamics of Broca’s area may likely play an important role in the encoding of speech sequences. However, electrocorticographic (ECoG) recordings with electrodes positioned over the surface of Broca’s area have shown that high-gamma activity increases prior to speech onset, but not during the time segment of speech production^13,41^, suggesting that Broca’s area underlies the preparation of the articulatory plan to be executed by the motor cortex but seems silent during the course of speech production. By contrast, an increased high-gamma power at some electrode sites positioned over the inferior frontal gyrus has been reported during the two phases of a speech perception to production task^42^. These seemingly contradictory findings can be reconciled by carefully dissociating *pars opercularis* from *pars triangularis*, as high-gamma activity has been observed to increase in *pars opercularis* but not in *pars triangularis* during the production of isolated words^43^. Given that high-gamma activity reflects the aggregated firing rate of the neural population captured by an electrode^44^, these results suggest that the firing rate in *pars triangularis* does not increase during time segments of speech production. Consequently, a reasonable hypothesis is that speech encoding in this specific region of Broca’s area could rely on fine changes in local neural dynamics while maintaining an unchanged overall spiking level. However, while large-scale single unit recordings are being increasingly explored during speech perception^45^ and production^46,47^, there has been to date no report of such recording in *pars triangularis* during speech production. It thus remains unknown whether and how the detailed temporal dynamics of spiking activity in this region, beyond its unchanged overall global rate, finely vary to encode produced speech.

Here, we investigated intracortical spiking dynamics in *pars triangularis* during overt and covert speech production in a patient undergoing awake surgery for the removal of a focal tumor located below Broca’s area. We confirmed that the overall amount of single unit spiking activity averaged over the array did not exhibit significant changes during overt or covert production of short sentences as compared to silence periods. However, several single units variably modulated their firing rates across conditions, some increasing and others decreasing depending on the cells and conditions. As a result, the overall rate-based population dynamics differed between overt and covert speech conditions. Moreover, we observed across the population of recorded cells that the temporal arrangement of spikes changed according to the sentences that were overtly pronounced. In particular, we found that, with an overall spiking level remaining constant, the neural population effectively encodes the content of uttered sentences through the temporal rearrangement of its spikes. We finally report that, contrary to individual spike rates, this spike-timing encoding remained largely conserved during covert speech imagination.

## RESULTS

A 29-year-old male patient was hospitalized at the Grenoble University Hospital for the removal of a focal left opercular low-grade glioma located below Broca’s area (Figure 1a). This resection was planned under awake surgery to limit possible post-operative functional damages to language functions. The surgical access to the tumor required the resection of the dorsal part of *pars triangularis*. This clinical indication thus offered an ethically justified opportunity to perform invasive intracortical recordings in this region before its resection. For this purpose, a 96-electrode Utah array was implanted anterior to the ascending branch of the inferior frontal gyrus (Figure 1b) while the patient was awake and recordings could be performed for about 20 minutes before the surgery had to resume. During this time, a list of 31 short French sentences or vowel sequences (see Supplementary Table 1) were prompted on a screen. For each of them, the patient was asked to read it once aloud, then to repeat it while it was no longer displayed on the screen, and finally to imagine repeating it again without any articulation (see Figure 1c-top and Supplementary Figure S1). The spiking activity of a total of 64 neurons could be isolated (see Methods and Supplementary Figure S2) to assess the ensemble dynamics of the population along the course of each trial (Figure 1c). We found that the overall rate of spiking of the whole population remained similar across the different read, repeat, covert and rest intervals of the trials (Figure 1d).

**Figure 1.**
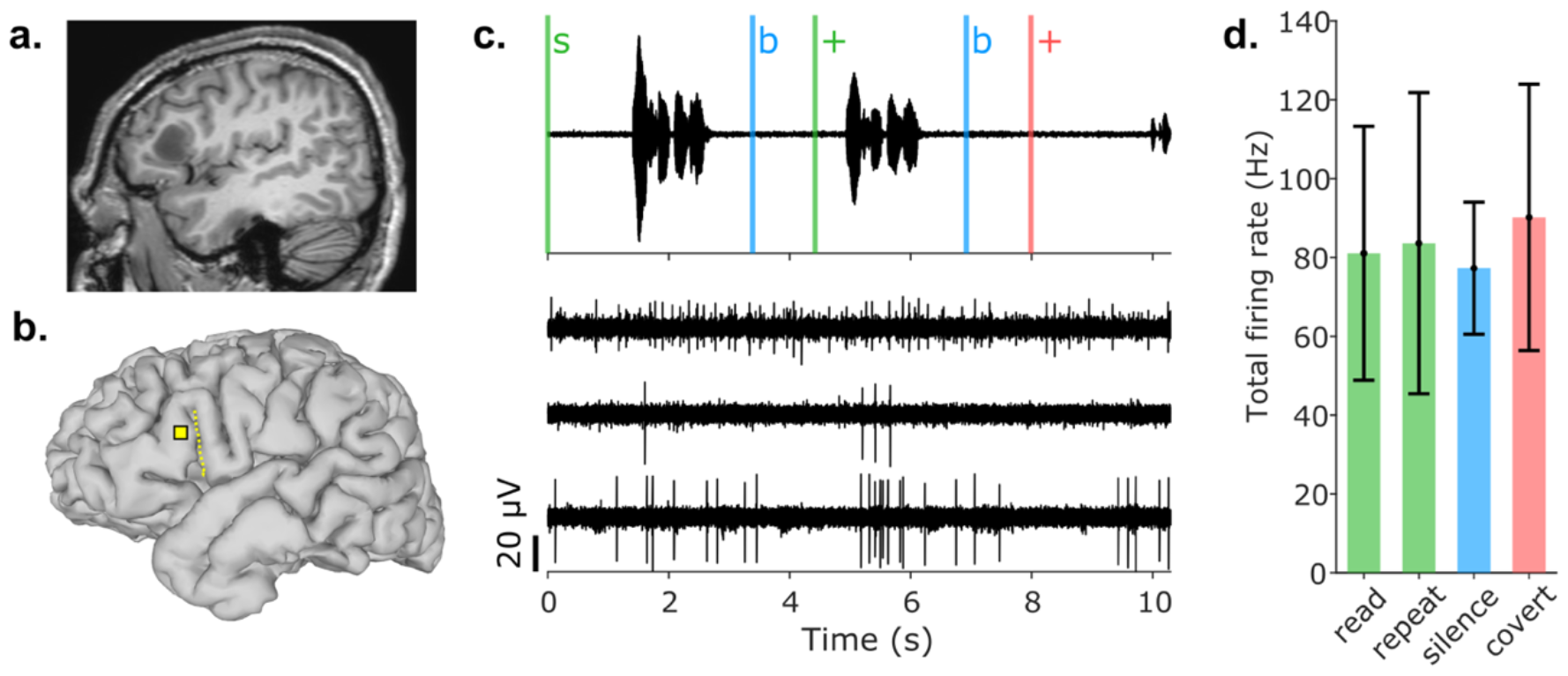
Global spike rate in a neural population of pars triangularis of Broca’s area during overt and covert speech. **a**. Preoperative MRI showing a focal tumor below Broca’s area. **b**. Localization of the Utah array with respect to the preoperative brain anatomy (dotted line indicates the ascending branch of the inferior frontal gyrus). **c**. Audio recording of a trial with onsets of visual cues displayed as vertical bars (top) and simultaneously recorded signals from 3 channels of the Utah array (bottom). The participant was instructed to read the sentence when it was displayed on the screen (s). After a blank screen (b), the participant was asked to repeat the sentence upon the appearance of the fixation cross (+). He finally had to covertly produce the same sentence after the appearance of a second fixation cross (also preceded by a blank screen). He indicated the completion of the covert repetition by saying aloud “OK”. All trials followed the same steps, with varying timings depending on the duration of the task completions and the manual triggering of cues “s” and “b” by the experimenter. The displayed neural signals were recorded from channels 57, 27 and 12 and band-pass filtered (see methods). **d**. Total firing rate gathering all units during the different read, repeat, silence and covert phases, averaged across all trials. The phases were labeled from the onsets of visual cues and speech production, as detailed in Supplementary Figure S1.

Next, we examined the spike rate of each single unit separately to determine whether this unchanged overall population firing was also observed at the level of individual neurons. The average firing rates of individual units over all considered trials ranged from 0.02 Hz for the slowest-firing unit to 7.4 Hz for the most active one, with an average of 1.3 Hz across units. For all units except one (unit 8801), no statistically significant difference in individual firing rates was observed between the overt reading and overt repetition conditions (either Kruskal–Wallis one-way rank analysis of variance across single unit firing rates in read, repeat, silence and covert phases showed *p* > 0.05, or, if not, post-hoc Bonferroni-adjusted Wilcoxon rank-sum test between the read and repeat conditions showed *p* > 0.05). To further investigate population dynamics across conditions, we thus opted to combine the read and repeat epochs into a single overt condition. Out of the 64 units, only 9 exhibited modulations of their firing rates across the three conditions of overt speech, covert speech, and silence. The nature of these modulations varied across units, with some units showing increased firing during overt speech production, others during covert speech imagination, and still others during the silence intervals preceding overt and covert speech production (Figure 2a). This finding suggests that while the overall firing rate of the global population remained at the same level, there were distinct variations in fine rate-based dynamics within a subset of cells across all three conditions. To further test this hypothesis, we computed the smoothed firing rates (see Methods) of the 9 modulated neurons and employed principal component analysis (PCA) to project the ensemble activity of this subpopulation onto a reduced 3D state space (Figure 2b and Supplementary Figure S3). We considered the first 4 principal components (PCs), which altogether represented 57% of the variance of the data. The first PC (20% of the variance) was not found to discriminate between conditions (Supplementary Figure S3a) but rather reflected the phases of preparation preceding overt and covert repetitions (Supplementary Figure S4). By contrast, within the subspace constituted by the next 3 PCs (37% variance), the population trajectory along the course of a trial was similar between the overt reading and overt repetition time intervals, but differed from that spanned during covert intervals (Figure 2b and Supplementary Figures S3b-c). The part of this subspace spanned by the population activity was indeed statistically different between the overt, covert and silence conditions (one-way multivariate ANOVA, p < 0.001; post-hoc pairwise Hotelling’s T^2^, Bonferroni-adjusted *p* < 0.001). This result thus confirms that, within a particular subspace of its firing rate, this population of *pars triangularis* neurons was differently activated between overt speech, covert speech and rest.

**Figure 2.**
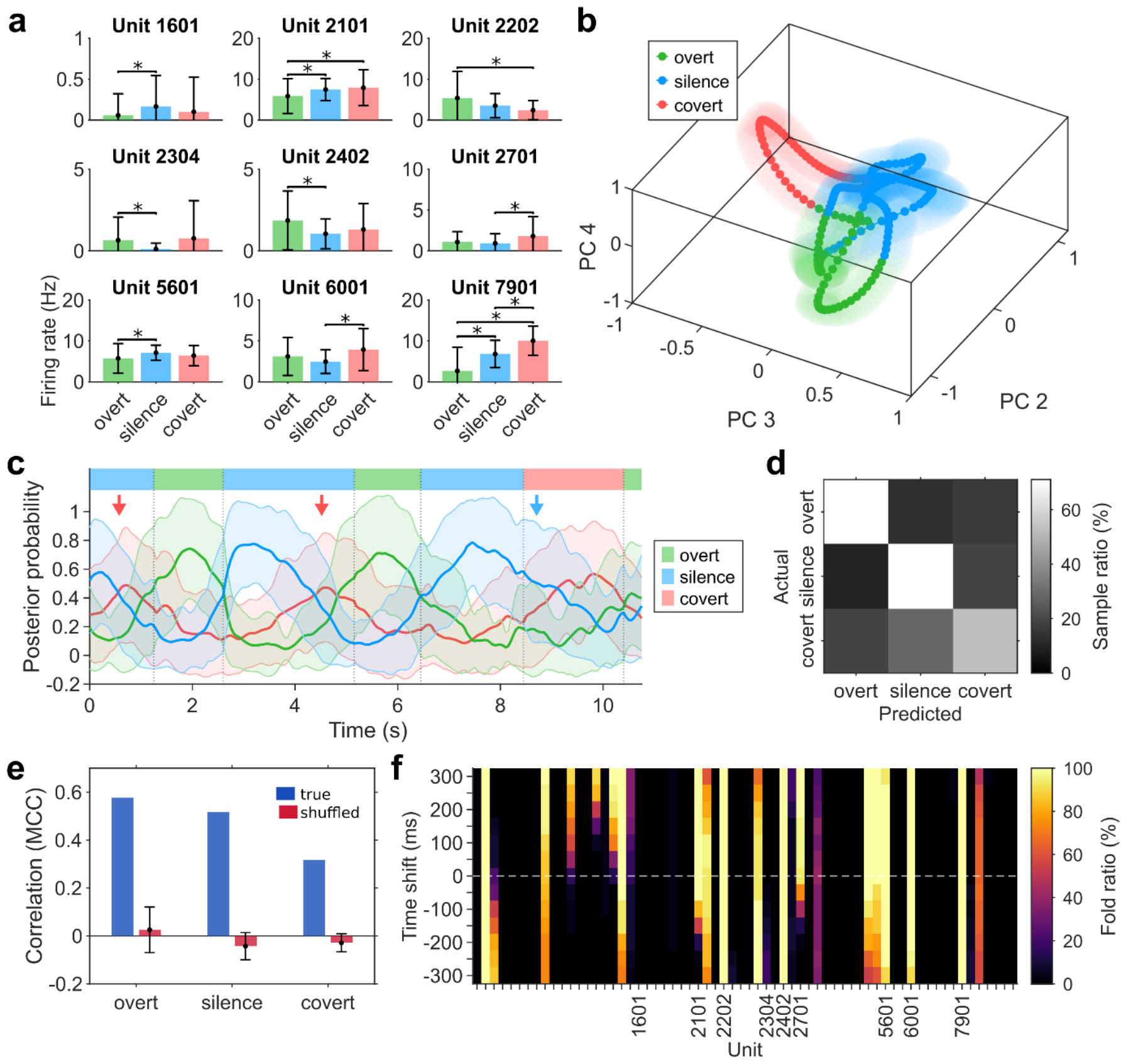
Population dynamics across overt speech, silence and covert speech intervals. **a**. Nine units were modulated across overt speech, silence or covert speech intervals. The average firing rates were computed in the labeled epochs (defined in Supplementary Figure S1). Units that produced different firing rates depending on the condition were identified using Kruskal–Wallis one-way analysis of variance (p < 0.05). Post-hoc tests were then computed to assess the modulation between each pair of conditions (Wilcoxon rank-sum test, Bonferroni-adjusted p < 0.05). **b**. Trial-averaged PCA projection of the firing rates of the 9 condition-modulated units shown in panel a. The components 2-4 produced a clearly visible separation of the 3 conditions (see Supplementary Figures S3 and S4 for other projections and the modulation of the 1st PC, respectively). The projected trajectories were averaged while preserving the different phases of the trial (see Supplementary Figure S1). The shaded areas are semi-transparent spheres whose radius is equal to the standard error of the mean (SEM). SEM was chosen instead of standard deviation in order to improve readability. **c**. Posterior probabilities of LDA classifier, averaged over all trials. Before computing the average, the posterior probabilities were computed on the whole trials using the classifiers trained on the labeled data (see Supplementary Figure S1). The solid lines and shaded areas indicate the mean and standard deviation of the posterior probability for each class. The colored patches at the top indicate the average phases of overt speech, silence and covert speech. The red arrows show two intervals of the trials where, on average, a short period of covert speech is predicted preceding overt speech production. The blue arrow indicates a predicted period of silence preceding the covert speech production that extends into this covert interval. **d**. Row-normalized confusion matrix of classification results showing the percentage of actual class samples predicted as samples of each of the three classes, for labeled segments only (see Supplementary Figure 1). Diagonal values are the true positive rates of overt, silence and covert and are equal to 70%, 71% and 52%, respectively. **e**. Evaluation of the performance of the classification for each of the 3 classes using Mathews correlation coefficient (see Methods). The chance levels were obtained on 20 random shuffling of the class labels. **f**. Selection ratio of the classification features. For each feature, represented by a unit and a time shift, the ratio of the number of selection occurrences over the 31 folds of the leave-one-out classification process is color-coded.

In order to reinforce this finding and determine if this different population activation could be seen at the single trial level, we further built a linear discriminant analysis (LDA) classifier to predict to which condition each time point of each trial belonged from the smoothed firing rates of the 64 units (see Methods). The classifier was trained and tested in a trial-wise leave-one-out fashion, using labeled segments (Supplementary Figure S1). During each fold, feature selection was computed on the training set and only features showing a statistically significant difference between at least 2 of the 3 classes were selected. As illustrated in Figure 2c, on average, the classifier predicted that the participant was performing overt speech during the two overt speech intervals, covert speech during the covert speech interval, and silence during the intervals preceding the beginning of each speech production. Interestingly, short covert speech intervals were predicted at the end of the silence intervals just before each overt speech production started (Figure 2c, red arrows). Also, the predicted period of silence preceding the covert speech production extended into this covert interval (blue arrow). Overall, the time points of the intervals labeled as covert were mostly predicted as belonging to covert intervals (53%) but also substantially as silence (29%), while overt and silence intervals were correctly predicted with accuracies over 70% (Figure 2d), and all conditions were predicted well above chance level (Figure 2e). These results thus indicate that *pars triangularis* is activated differently during overt speech, covert speech, and silence at the single trial level. From the average feature selection over the different folds (Figure 2f), we observed that only a subpart of all smoothed firing rates consistently showed a statistically significant difference between at least one of the two classes. It also appears that values of the smoothed firing rates following the sample to be classified (positive time shifts) were more often selected than preceding activity.

We then questioned whether *pars triangularis* also differently encoded the different individual speech sequences that were overtly produced. To this end, for all utterances, we extracted the spike trains on a 2-second window starting at speech onset. In several cases, we observed a striking similarity of the spike sequences produced by subsets of neurons during two utterances of the same sentence (Figure 3a). To further quantify this observation, we computed, for each unit and each pair of produced sentences, two complementary and uncorrelated measures of spike train similarity: one sensitive to the difference in the number of spikes (*d*_*count*_) and the other in the timing of spikes (*d*_*timing*_) between both trains. The distance *d*_*timing*_ was derived from the spike train distance proposed by Victor and Purpura^48,49^ but normalized so as to be uncorrelated with spike counts (see Methods). This timing-based distance can be assimilated to the cost of transforming the sequence of inter-spike intervals (ISIs) of one train into that of the other train. It integrates a temporal precision parameter *t*_*p*_ representing the maximum difference in duration that is allowed to consider that two ISIs are equivalent. In other words, the value of *t*_*p*_ can be interpreted as the temporal jitter of spike times allowed to consider the two trains as similar. The two distances between spike trains computed for each individual unit were then averaged across units to form population-wide spike train distances *D*_*count*_ and *D*_*timing*_ (see Methods). These two population-wide distances were computed for each pair of utterances, to measure the similarity of their underlying neural activity. In the case of *D*_*timing*_, the procedure was repeated for different values of *t*_*p*_ ranging from 1 millisecond to 100 seconds, in order to determine whether a particular precision of spike intervals specifically reflected any encoding of the sentences.

**Figure 3.**
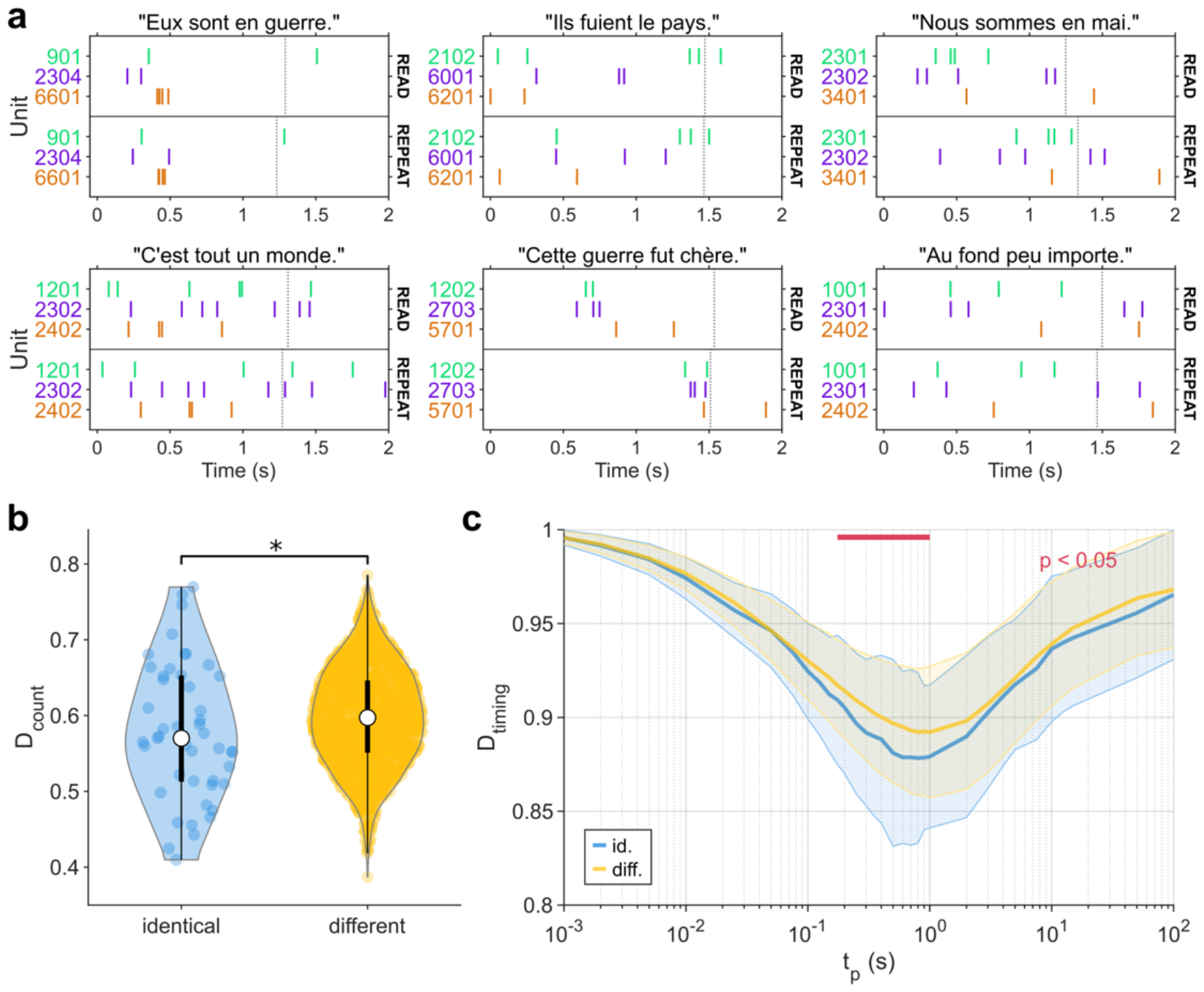
Population-wide spike train distances during the production of identical vs. different sentences. **a**. Examples of spike trains occurring during two overt repetitions of the same sentence (displayed above the graphs). In each graph, the raster plots show the activity of 3 units. The x-axis represents the time from speech onset. The vertical dashed lines indicate speech offsets. **b**. Distributions of the population-wide spike train distance D_count_ computed for pairs of either identical (blue) or different (yellow) sentences. Values of D_count_ for identical sentences were statistically significantly lower than for different sentences (one-tailed Mann-Whitney U-test, n_1_=47, n_2_=1844, z=-1.71, p=0.044). **c**. Distributions of the population-wide spike train distance D_timing_ for either identical (blue) or different (yellow) sentences as a function of the temporal precision t_p_. Solid lines represent medians and shaded areas cover ±1 MAD-derived standard deviation (see Methods).

We first tested whether these population-wide distances were lower for identical versus different uttered sentences. We found that values of *D*_*count*_ were indeed lower for identical sentences than for different sentences (Figure 3b), indicating that the number of spikes produced by individual units were more similar for identical vs. different sentences. At the same time, values of *D*_*timing*_ were found to be lower for identical vs. different sentences for spiking temporal precisions *t*_*p*_ between 175 and 1000 ms (Figure 3c), indicating that the temporal spiking pattern within a precision of a few hundreds of milliseconds also characterized which sentence had been produced.

To further confirm that both the number and the temporal arrangement of spikes encode overtly produced speech in *pars triangularis*, we then questioned whether each population-wide distance could be used as the decision criterion of a template matching classifier to decode which sentence has been pronounced by the participant. This decoding scheme attributed to each population spike train generated during the production of a sentence, the sentence label corresponding to its lowest population-wide distance to any other sentence (see Methods). As previously, this procedure was tested with different temporal precision *t*_*p*_ ranging from 1 millisecond to 100 seconds to determine the precision of spike intervals reflecting the encoding of the sentences. When using *D*_*timing*_, the classification accuracy was found to be statistically significantly above the one obtained after temporally shuffling spike instants for a wide range of *t*_*p*_ values, and maximal for temporal precisions of 250 and 300 ms (Figure 4a, solid blue curve). In that case, about 29% of the sentences could be assigned the correct label, which was significantly higher than for surrogate data obtained by shuffling either the sentence labels (empirical *p* < 10^-3^) or the times of the spikes within trains (empirical *p* < 10^-3^), which lead to chance decoding levels of at most 8% and 10%, respectively (Figure 4a, dotted grey curve and dashed orange curve, respectively). When using *D*_*count*_, the decoding accuracy was slightly higher at 31%, and also statistically significantly well above that obtained after shuffling the sentence labels (empirical *p* < 10^-3^). Because two identical overt sentences were always produced one after the other in a given trial, we checked that the decoding performance was not due to the proximity in time between the read and repeat intervals. To this end, the same decoding procedure was repeated on time-shifted read and repeat windows. For each trial the shift was defined in order to make the end of the surrogate read window coincide with the sentence appearance. The surrogate read and repeat windows were separated by the same amount of time as the original windows. In this case, the decoding performance was found to be only slightly over chance level (see Supplementary Figure S5).

**Figure 4.**
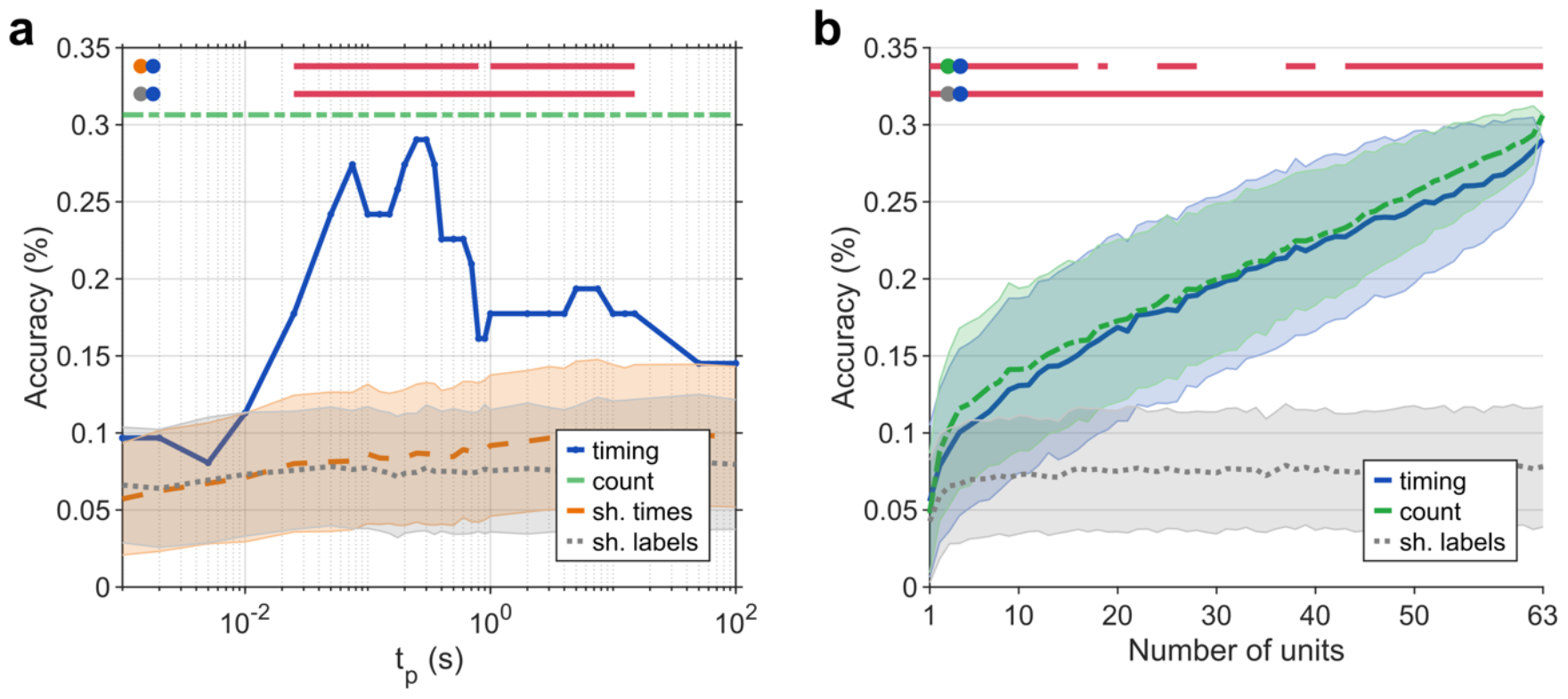
*Classification of overtly produced sentences from ensemble spiking activity recorded in* pars triangularis. ***a***. *Decoding accuracy based on D*_*timing*_ *for different values of temporal precision t*_*p*_, *using the original data (solid blue curve), shuffled spike trains (dashed orange curve) and shuffled sentence labels (dotted grey curve). For comparison, the level of accuracy obtained using count-based distances is indicated by the horizontal green line. The horizontal red lines indicate the ranges of t*_*p*_ *values for which the timing-based decoding accuracy is statistically significantly higher (empirical p < 0*.*05) than for the surrogate data (top: surrogate=shuffled spike times, bottom: surrogate=shuffled sentence labels)*. ***b***. *Decoding accuracy based on D*_*count*_ *and D*_*timing*_ *as a function of the number of randomly selected units. The classification based on D*_*timing*_ *uses t*_*p*_ *= 300 ms. The chance level is computed by applying the classification based on D*_*timing*_ *to datasets with shuffled sentence labels. The horizontal red lines indicate the ranges of numbers of units for which D*_*timing*_ *accuracies are significantly different (p < 0*.*05, two-tailed Wilcoxon ranksum test, n*_*1*_*=1000, n*_*2*_*=1000) from D*_*count*_ *accuracies (top) or shuffled labels accuracies (bottom). All cases of shuffling consisted of 1000 repetitions. All distributions are represented by their means and standard deviations*.

Previous studies aiming at decoding neural population activity based on individual firing rates have consistently shown that decoding accuracy strongly depends on the number of recorded neurons ^50–52^. We thus tested the influence of the number of units used for the decoding approach on its accuracy, for both count-based and timing-based distances. As shown in Figure 4b, we found that in both cases the average classification accuracy consistently increased with the number of considered units, and was statistically significantly higher than the accuracy obtained with shuffled sentence labels for any (*D*_*timing*_) or most (*D*_*count*_) number of units. Moreover, for any number of units, the accuracy obtained using *D*_*count*_ was always equivalent or slightly higher than when using *D*_*timing*_ with *t*_*p*_ = 300 ms (Figure 4b). The distances based on spike counts and timing being uncorrelated (See Supplementary Figure S7), these results thus indicate that both the number and the timing of spikes encode overtly produced speech in *pars triangularis*.

Finally, we questioned whether the precise timing of the spikes characterizing the overt pronunciation of a sentence was preserved during the covert repetition of the same sentence. To this end, we conducted the same template matching classification scheme, but this time attributing to each covert population spike train the sentence label that corresponded to its smallest distance to the different overt repeat population spike trains (see Methods). The distance criterion was either *D*_*timing*_ or *D*_*count*_. The onset of covert speech being unknown, we considered that the covert speech onset occurred 600 ms after the fixation cross appearance, which corresponded to the average speech onset delay in the repeat condition (see Methods). As shown in Figure 5a (solid blue curve), we found that covertly produced sentences could be classified based on *D*_*timing*_ with an accuracy up to 23%, this maximum being obtained with temporal precisions *t*_*p*_ of 300 and 350 ms. This timing-based decoding accuracy was statistically significantly above the surrogate accuracies obtained after randomly shuffling the spike times within trains (empirical *p* < 10^-3^) or after shuffling sentence labels (empirical *p* < 10^-3^). When using *D*_*count*_, the decoding accuracy was only 13% (Figure 5a, dashed green line), yet still statistically significantly above that obtained after shuffling the sentence labels (empirical *p* < 0.015). Thus, the timing-based decoding accuracy was 75% higher than the count-based decoding accuracy when matching covert population spike trains onto overt repeat population spike trains. Finally, we compared decoding accuracies based on *D*_*count*_ and *D*_*timing*_ for different numbers of units used for the classification process. Timing-based accuracies (with *t*_*p*_ = 300 ms) were above chance level for any number of units and higher than count-based accuracies for almost any number of units (Figure 5b). Altogether, these results confirmed that the spike-timing encoding of overtly produced sentences was largely conserved during covert speech, more than rate-based encoding, a phenomenon illustrated for a few examples of sentences in Figure 5c.

**Figure 5.**
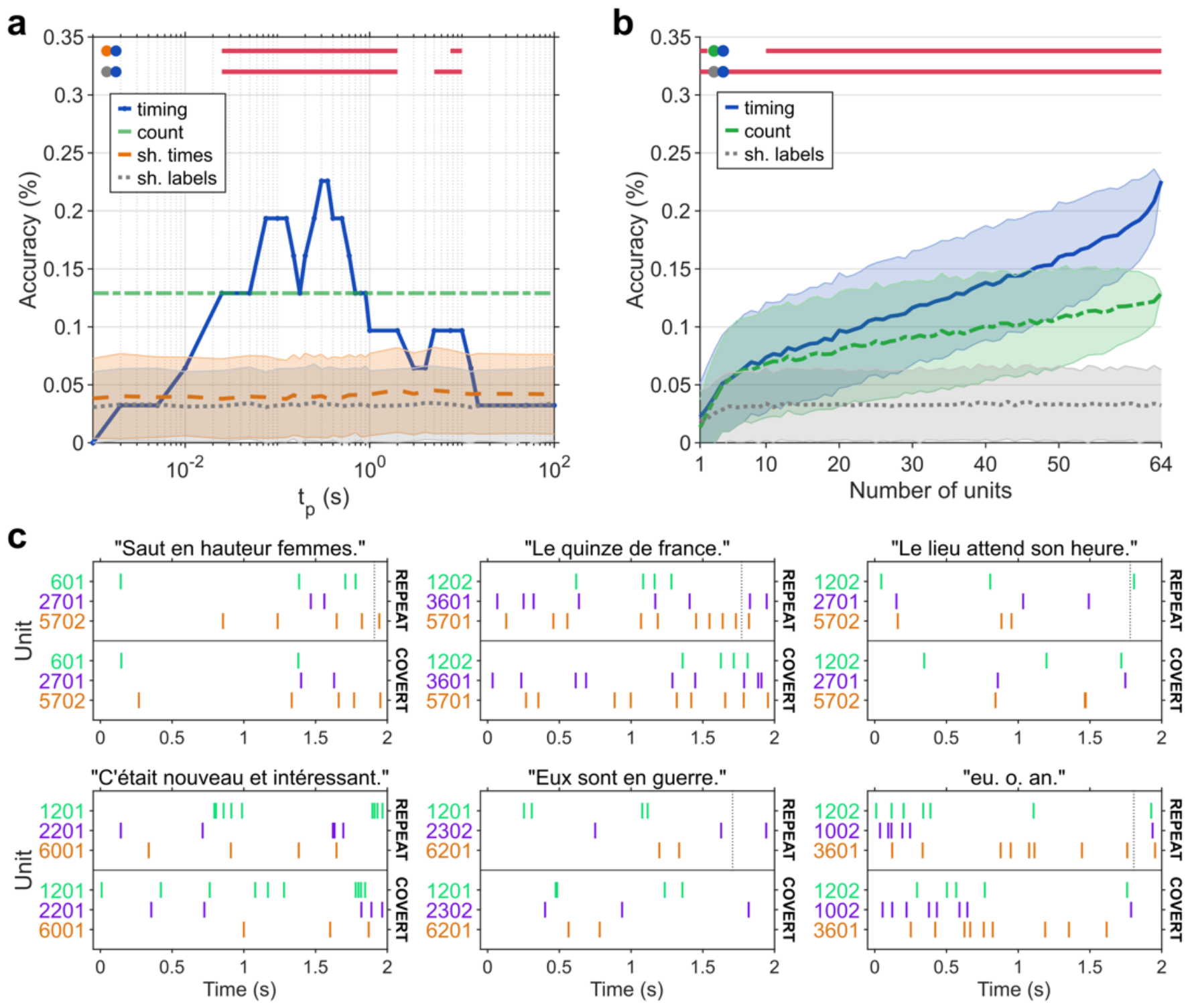
*Classification of covertly produced sentences by matching population spiking activity recorded in* pars triangularis *during covert repetitions to that recorded during overt repetitions*. ***a***. *Decoding accuracy based on D*_*timing*_, *for different values of temporal precision t*_*p*_, *using the original data (solid blue curve), shuffled spike trains (dashed orange curve) and shuffled sentence labels (dotted grey curve). For comparison, the level of accuracy obtained using count-based distances is indicated by the horizontal green line. The horizontal red lines indicate the ranges of t*_*p*_ *values for which the timing-based decoding accuracy is statistically significantly higher (empirical p < 0*.*05) than for the surrogate data (top: surrogate=shuffled spike times, bottom: surrogate=shuffled sentence labels)*. ***b***. *Decoding accuracy based on D*_*count*_ *and D*_*timing*_ *as a function of the number of randomly selected units. The classification based on D*_*timing*_ *uses t*_*p*_ *= 300 ms. The chance level is computed by applying the classification based on D*_*timing*_ *to datasets with shuffled sentence labels. The horizontal red lines indicate the ranges of numbers of units for which D*_*timing*_ *accuracies are significantly different (p < 0*.*05, two-tailed Wilcoxon ranksum test, n*_*1*_*=1000, n*_*2*_*=1000) from D*_*count*_ *accuracies (top) or shuffled labels accuracies (bottom). All cases of shuffling consisted of 1000 repetitions. All distributions are represented by their means and standard deviations*. ***c***. *Examples of spike trains occurring during overt and covert repetitions of the same sentence (displayed above the graphs). In each graph, the raster plots show the activity of 3 units. The x-axis represents the time from overt speech onset and estimated covert speech onset. The vertical dashed lines indicate overt speech offset*.

## DISCUSSION

Broca’s area has been identified as a key structure of overt and covert speech production using imaging techniques, but its population dynamics has remained largely unexplored. How speech is encoded at the population level and whether this encoding differs between overt and covert speech remain two unresolved questions. This study aims to address these interrogations by offering initial answers on population-level speech encoding in *pars triangularis*.

First, we found that the overall spike rate of the population remained unchanged during both overt and covert speech periods as compared to silence periods. This result is in accordance with previous fMRI^25^ and ECoG^13^ studies showing that, compared to *pars opercularis, pars triangularis* does not feature strong activation for simple reading and repetition tasks. It also corroborates previous reports revealing no particular increase in high-gamma ECoG activity (reflecting population spiking^44^) in Broca’s area, and thus its apparent silence^41^, during spoken responses, while consistent activity increase is observed over the motor cortex^41,53^. This is also in line with a recent rate-based speech brain-computer interface study reporting highly efficient speech reconstruction from intracortical motor cortex spiking activity but a poor contribution of Broca’s area in such speech reconstruction^46^. Moreover, within this unchanged overall population rate, we observed that only a small proportion of neurons (14%) modulated their firing individually across the different states (silence, overt, and covert speech). These first results suggest that the modulation of individual spike rates may likely not be the key mechanism by which pars triangularis encodes produced speech, leaving as another encoding alternative the possibility that the population rearranges its fine temporal dynamics at the level of individual neural firings.

The second main finding of the present work confirms in more details this initial suggestion. We indeed found that the temporal arrangement of spikes emitted by subsets of neurons was specific to particular sentences. Typically, action potentials of a few cells were emitted similarly in time for different occurrences of a given sentence, while this occurred for another subset of units for a different sentence (Figure 3a). This phenomenon was robust enough to be used to decode which sentence was produced with an accuracy very significantly above chance level, and comparable to that obtained from spike counts, despite the low number of trials allowed by the awake surgery recording condition (Figure 4a-b). This finding thus suggests that, in Broca’s area, neural assemblies^54,55^ within *pars triangularis* may encode dynamically the flow of speech that is being produced through the specific temporal arrangement of their ongoing flow of spikes.

As a consequence, the present results also propose a new type of feature for speech decoding. Classical schemes of neural encoding of motor behavior are indeed based on spike rates, the modulation of which is a hallmark of primary motor neurons engaged in the realization of a movement^56^. As a result, single and multi-unit activity rates have been the main neural features used for motor movement decoding and intracortical BCIs based on the activity recorded over the motor cortex^50,57–64^. Further studies have also shown successful speech decoding from single/multi-unit spike rates^65–71^ or high-gamma amplitude, an indirect signature of multi-unit activity rates^44^, recorded over the sensorimotor cortex or distributed depth electrodes^72–76^. Here, we found that the accuracy of overt speech decoding in *pars triangularis* was comparably efficient when based on the similarity of population spike timings than when based on the similarity of spike counts (Figure 4). Moreover, we found that spike timing could be successfully used to predict covertly produced sentences by matching their corresponding spike trains to those occurring during overt speech. In particular, spike-timing-based decoding allowed a 75% higher decoding accuracy of covert sentences than when using spike counts (Figure 5). This superiority of timing-based decoding might be explained by the fact that individual firing rates differed between overt and covert speech (Figure 2), which likely prevented a strong decoding of covert sentences from overt rate codes. By contrast, despite the fluctuations of individual spike rates, the spike-timing similarity was more robustly conserved between overt and covert speech. Moreover, the temporal precision of spiking that enabled the best sentence decoding was also found to be the same (300 ms) between overt and covert speech, indicating that a similar spiking precision of 300 ms underlies the encoding of overt and covert sentences in *pars triangularis*. Altogether, these results support the existence of a spike-timing code for overt speech in this region that remains largely conserved during speech imagination, which may open new perspectives for the development of speech BCIs based on single-unit recordings in Broca’s area.

Theoretical neural computation studies have proposed that spike rates are likely not the most efficient way to encode behavior and that more frugal and reactive neural codes based on precise spike timing and synchrony could be more efficient^77–79^. This hypothesis has been supported by several experimental studies confirming that coordinated firing constitutes an alternative neural code to pure spike rates. As a few examples: encoding of behavior without modulation of firing rates has been reported in frontal premotor cortex^80^, small subsets of neurons in the primary motor cortex of non-human primates have been shown to synchronize firing for certain directions of movements during a center-out task, different subsets being concerned for different directions of movements^81^, and encoding of sensory information by spike timing has been observed in the auditory cortex, the retina and the somatosensory cortex^82–86^. To our knowledge, however, we present here the first evidence for cortical spike timing encoding of speech in humans.

This phenomenon might generalize sequence-based coding of simpler vocal behavior previously observed in non-human species. Precise spike timing has been shown to encode song production in the vocal cortex of Zebra finches. In this species, the emission of a song is underlain by trains of short bursts emitted by neurons of the RA motor area and highly precise sequences of very sparse spikes emitted by RA-projecting neurons of the HVC premotor area^87,88^. Focal cooling of the HVC area induces a temporal stretching of produced songs^89^. Similarly, in humans, focal cooling of Broca’s area including *pars triangularis* alters the temporal structure of produced speech^40^. Typically, each HVC_RA_ neuron emits at most one burst during a song motif^88,90^ and receives stereotyped trains of synaptic inputs at multiple time points along the song motif that are reproducible across different renditions of the same motif ^90,91^. These temporal sequences have been attributed to an intrinsic chain-like wiring of HVC^90^. While such model is well adapted to the songbird repertoire composed of a single stereotyped song, it might however lack versatility to encode the nearly indefinite combinatorial speech repertoire of humans, as it would require a nearly infinite number of predefined chained subnetworks. A possible alternative suggested by the present observations, could be the existence, in humans, of a network encoding dynamically the variety of possible speech sentences through timely precise and distributed firing of a cortical neural population that involves *pars triangularis*.

## ACKNOWLEDGEMENTS

We thank participant P3 for his involvement in the experiment during awake surgery, as well as Rob Franklin and Sherman Wiebe from Blackrock Microsystems for their assistance. This work was supported by the FRM Foundation under Grant No. DBS20140930785, the French National Research Agency under Grant Agreement No. ANR-16-CE19-0005-01 (Brainspeak) and in the framework of the “Investissements d’avenir” program (ANR-15-IDEX-02), and by the European Union’s Horizon 2020 Research and Innovation Program under Grant Agreement No. 732032 (BrainCom).

## AUTHOR CONTRIBUTIONS

FB, SC and BY prepared and performed the recording. FB developed software codes for data acquisition and processing. SC implanted the electrode array during awake surgery. PR and BY designed the analysis of the data. PR performed the analyses. BY and PR wrote the paper. BY designed and supervised the study and obtained appropriate funding.

## METHODS

### EXPERIMENTAL PROCEDURE

#### Human participant

The electrophysiological recording on which this study is based was acquired from a 29-year-old participant, P3, during his awake surgery for tumor resection at the Grenoble-Alpes University Hospital (*Centre Hospitalier Universitaire Grenoble-Alpes*). This recording was performed with the informed consent of the participant, within the Brainspeak clinical trial (NCT02783391) approved by the French regulatory agency ANSM (DMDPT-TECH/MM/2015-A00108-41) and the local ethical committee (CPP-15-CHUG-12).

#### Experimental task

Data was acquired in successive trials. In each trial, the participant was asked to read, repeat and covertly repeat a French sentence displayed on a screen. The trials were driven by cues displayed on a screen as shown in Figure 1. Each phase of the trial was started manually in order to adjust to the condition of the patient during the awake surgery. On average, the read, repeat and covert phases lasted 1.47 ± 0.31 s, 1.44 ± 0.31 s, and 1.94 ± 0.30 s, respectively. A trial started with the sentence prompted on the screen until the patient completed its reading. Then, a black screen was prompted for 1 sec at the beginning of the repeat phase, after which a cross remained on the screen for the duration of the phase until the patient completed repeating the sentence. Finally, a black screen was prompted for 1 sec at the beginning of the covert phase, after which a cross remained on the screen for the duration of the phase until the patient pronounced “OK” to indicate that he has completed the covert imagination of the sentence. Out of the 43 recorded trials, only the first 31 were analyzed, for reasons exposed in the following. The chosen sentences were part of the articulatory-acoustic corpus BY2014^92^. The corpus of sentences in the experiment contained a majority of verbal sentences, some nominal sentences and four sequences of vowels (for example “a, i, ou”). Each vowel sequence was used in two trials, all other sentences were used only in a single trial. The 31 selected trials contain 27 different sentences, 62 speech epochs and 31 covert speech epochs.

#### Electrophysiological recording

Brain activity from participant P3 was recorded in the operating room prior to tumor resection. A 96-channel intracortical Utah microelectrode array (UEA, Blackrock Microsystems, USA) with 1.5-mm long electrodes was inserted in *pars triangularis* of Broca’s area (Figure 1), at a location that was subsequently resected to access the tumor for its removal. The pedestal, serving as ground, was screwed to the skull. Two wires with deinsulated tips were inserted below the dura, and one was used as reference. The electrodes were connected via a Patient Cable (Blackrock Microsystems, USA) to a front-end amplifier (Blackrock Microsystems, USA) where signals were digitized at 30 kHz, and further transmitted through an optic fiber to a Neural Signal Processors (NSP, Blackrock Microsystems, USA). The signals were filtered with the integrated digital filter of the NSP between 0.3 and 7500 Hz and transmitted to a computer for storage.

#### Audio recording

The participant’s speech was recorded along with his neural data. A microphone (SHURE Beta 58 A) was positioned at about 10-20 cm from his mouth. The signal was amplified using a microphone preamplifier (Roland OCTA-CAPTURE) and digitized by the NSP, at the same rate and synchronously with the neural data.

### SEGMENTING AND LABELING TRIALS

#### Trial phases

Each trial was broken down into different phases using manually-annotated start and end times of utterances as well as recorded trigger times corresponding to stimulus display (Supplementary Figure S1):

- “read”, “repeat” and “OK” overt speech phases, delimited using audio
- a “covert” phase, delimited by the instruction stimulus and the “OK”
- “silence” phases, in-between the previous phases

When averaging the results from different analyses across trials in Figures 2b-c and Supplementary Figure S4a, the boundaries between each phase were aligned so as to preserve the phase-related modulations. Before averaging, the data was transformed so that phases have the same duration in each trial. For each phase the average duration across trials was computed. Each trial was temporally stretched or compressed so that its duration matched the average duration. Signals were temporally warped using linear interpolation.

#### Phase labels

In order to classify firing rates according to conditions, labels corresponding to overt speech, silence and covert phases were defined in each trial (Supplementary Figure S1). The labeled epochs were defined as subparts of trial phases. The periods preceding overt speech were excluded in order not to risk to include neural activity related to speech preparation and stimulus presentation in the silence data. In order not to include samples for which labeling was ambiguous, 300-ms margins at the beginning and end of each phase were also excluded. At the beginning of covert speech epochs, a 600-ms margin was removed instead, in order to take into account the participant’s reaction time before actual covert speech. 600 ms indeed corresponds to the rounded value of the average reaction time in the overt repetition task. To evaluate the classification process, chance levels were estimated by randomly shuffling the labels of these labeled epochs.

### NEURAL DATA PROCESSING

#### Common average reference

The problem of common average reference (CAR) is that it can introduce channel-specific noise, in particular high-amplitude transient artifacts, to the other channels. In order to mitigate the propagation of channel-specific noise to all channels, the median value of all channels was used as a reference and subtracted. Using median instead of mean in common average reference has been shown to increase the number of detected spikes in intracortical micro-electrode recordings^93^.

#### Spike sorting

Prior to spike-sorting, band-pass filtering was applied to the neural data. The band-pass filter was designed to select frequency content between 400 and 3000 Hz. The standard deviation of each channel was estimated using the median absolute deviation, using the following formula where Φ is the reciprocal of the quantile function.

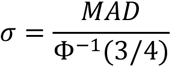

This method allows to mitigate the influence of high-amplitude artifacts and to get a close estimate of the standard deviation of the signals without spikes^94^. The neural signal of each channel was set to zero mean and scaled to have a standard deviation of one. The baseline standard deviation designates the estimation of the standard deviation from the median absolute deviation^94^. The spike sorting process was carried out using Offline Sorter (Plexon, USA). The spike detection thresholds were set to plus and minus 5 times the MAD-based standard deviation of each channel. Waveforms were extracted by selecting the 2-ms window around the detected peaks (0.7 ms before the peak and 1.3 ms after). The detected waveforms were projected in a feature space using PCA. Spike-sorting was then performed in a semi-automatic way. First, the T-Distribution expectation-maximization algorithm was applied. This algorithm fits a mixture of T-Distributions to the point densities in feature space by adjusting the number of distributions and their parameters. Then, waveforms and clusters were manually adjusted. 64 putative neurons (or units) were obtained from 31 channels. Each unit was designated using a unique 4-digit number where the 2 first digits refer to the channel on which they were recorded. Most of the units were not detected anymore after trial 31, probably due to a relative movement between the implant and the brain. For this reason, only the first 31 trials were analyzed in the following steps. An example of spiking data is illustrated in Supplementary Figure S2.

#### Smoothed firing rates

Smoothed firing rates were computed by convolving the spike trains with a Gaussian kernel^95^. This method, relying on the hypothesis of rate coding, was developed as an alternative to trial-averaged peri-stimulus time histograms (PSTHs). It is useful to study the activity of single neurons on single trials, especially when the activity is not precisely time-locked to external events. Here a Gaussian kernel with a standard deviation of 300 ms was chosen. This kernel is wide compared to the ones used in other studies, which is coherent with the comparatively low firing rates in our dataset.

### CONDITION CLASSIFICATION

#### Features

In order to assess the difference of firing rates between overt speech, silence and covert speech, classification of those 3 conditions was carried out. The performance of the classifier was assessed by leave-one-out cross-validation on the 31 trials. The smoothed firing rates sampled at 20 Hz were used as features for the classification. Each observation at a given time *t* was composed of the firing rates of the 64 units within a window centered around time *t*. The context covered the 300-ms periods preceding and following the observation time, resulting in a neural features space with 64 × 13 = 832 dimensions.

In each fold, a subset of features was first selected using a Kruskal-Wallis one-way analysis of variance. For each feature, this test returned the *p*-value for the null hypothesis that the feature’s distribution was identical for the 3 classes. Features with a *p*-value inferior to 0.01 were pre-selected. Then pairwise Wilcoxon rank-sum tests were used and only features with a Bonferroni corrected *p*-value inferior to 0.01 were selected.

#### Linear discriminant analysis

The classification used linear discriminant analysis (LDA). This model was chosen for its simplicity, as is has a no hyperparameters to tune and is not prone to overfitting due to its low number of internal parameters. LDA fits multivariate Gaussian densities to each class. It assumes that the feature data are normally distributed and that the covariance matrices of the classes’ Gaussian distributions are equal. During training, the mean of the Gaussian distributions and their common covariance matrix are estimated so as to minimize a misclassification cost.

When applied to an observation, the trained LDA model returns for each class the posterior probability that the observation belongs to it. This posterior probability is a product of the prior probability and the modeled multivariate normal density. Here the prior probability, the probability for an observation to belong to a class independently of its value, was considered uniform, that is to say equal to 1/3 for all 3 classes.

#### Mathews’ correlation coefficient

In addition to the confusion matrix, the classification results were evaluated in a more concise way using the Matthews correlation coefficient (MCC). The MCC is an evaluation metric for binary classification that is computed using the following formula:

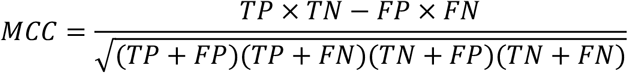

In this formula, TP is the number of true positives, TN the number of true negatives, FP the number of false positives and FN the number of false negatives. The MCC returns the same values as Pearson’s correlation coefficient computed between the actual and predicted labels. It returns +1 for a perfect classification and -1 if the prediction is the exact opposite of actual labels. MCC was chosen because it has advantages over the commonly used metrics of accuracy and F1-score, especially on imbalanced datasets^96^. It was used to evaluate the classification of each class against the two others.

### SPIKING ACTIVITY ANALYSIS

#### Spike trains and population spike trains

A spike train is defined as a list of successive spikes produced by a given unit on a given temporal window. Practically, it consists of the list of the time of occurrence of the extracted spikes relative to the beginning of the temporal window. A population spike train refers to the collection of spike trains produced by all considered units on a given temporal window. We note 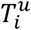 the spike train corresponding to unit *u* on window *i* and *T*_*i*_ the population spike train corresponding to the same window. The number of spikes in 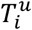 is noted 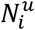.

#### Temporal shuffling of spike trains

Temporal shuffling of spike trains is used to create surrogate spike trains with preserved spike counts but different temporal patterns. A temporally shuffled version of a spike train is obtained by replacing the time of each spike by a value sampled from a uniform distribution on the interval extending from 0 to the duration of the window that was used to extract the spike train. The k^th^ shuffling of spike train 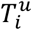 is noted 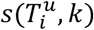.

#### Victor and Purpura (VP) spike train distance

In order to estimate the similarity between spike trains for a given unit, we initially considered the Victor and Purpura distance (VP distance) based on inter-spike intervals (ISIs)^48,49^. Given that repetitions of the same sentence are not necessarily produced at the same pace, the distance based on inter-spike intervals was preferred to the one based on spike times. Indeed, a global time shift applied to several successive spikes affect the timing of each spike but only the interval preceding the first spike. Moreover, a global stretching of an interval of spikes will affect each ISI by a little amount of time but will affect individual spike times increasingly along time.

This metric is computed as the cost of transforming a spike train 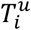 into another spike train 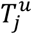. This transformation is accomplished through a succession of elementary steps whose individual costs are added together to form the total cost. The distance between 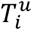 and 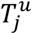 is given by the smallest total cost of all sequences of elementary steps that are needed to transform 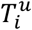 into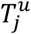 . The available elementary steps are: adding an interval, deleting an interval and changing the duration of an interval. Both adding and deleting an interval are assigned a cost of 1. The cost of changing the duration of an interval is controlled by the parameter *q* so that changing its duration by Δ*t* is assigned the cost *q*|Δ*t*|.

If *q* = 0, the distance between two spike trains is equal to the difference in the number of intervals (and thus to the difference in the number of spikes). If *q* tends towards infinity, the distance between two spike trains (which are not perfectly synchronized) reaches a limit value equal to the sum of the number of intervals in the two trains. For intermediate values, *q*, measured in s^-1^, expresses the sensitivity of the metric to the timing of spikes. When comparing two spike trains S_a_ and S_b_ containing only one interval each (two spikes), the two candidate sequences of elementary operations to transform S_a_ into S_b_ are either 1) changing the interval of S_a_ or 2) deleting and inserting an interval in S_a_. Changing the interval will result in a lower cost only if the two intervals differ by less than 2/*q*. Consequently, we decided to analyze the VP distance with respect to a parameter *t*_*p*_ = 2/*q*, representing the temporal precision considered to match trains. The VP distance between 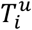 and 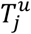 for temporal precision *t*_*p*_ is noted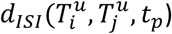.

When computing the distances between two overt sentences, we modified this distance to include the interval between the beginning of the window from which the spike train is extracted and the first spike. This allowed to include trains with single spikes and to consider that two spike trains are closer if they start with similar latency after speech onset than if not. This initial interval was discarded when comparing overt and covert speech activities as the covert speech onset was not precisely known. When there were not enough spikes to compute *d*_*ISI*_ this distance was considered as missing for this pair of trains and unit in further computations.

#### Count- and timing-sensitive spike train distances

By construction, the VP distance *d*_*ISI*_ is sensitive to the temporal arrangement of spikes but also to the number of spikes in the considered trains. This is illustrated in Supplementary Figures S6a and S6b showing that, on a temporally shuffled versions of the dataset, *d*_*ISI*_ is positively correlated to the sum of the spike counts of the two considered trains and to their difference, respectively. In order to decouple the influence of the number of spikes from that of the timing of the spikes, we defined two different normalized spike train distances: one designed to be sensitive to spike counts, and one to the temporal arrangement of spikes.

The normalized count-sensitive distance is defined as the absolute difference of spike counts divided by the total count:

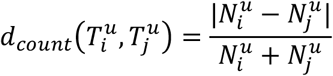

The normalized timing-sensitive distance is computed by dividing the value of *d*_*ISC*_ for two spike trains by its average value obtained for shuffled versions of the spike trains:

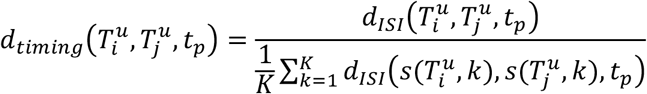

where *s()* is a shuffling operation on the time instants of all spikes of a given train (see above). The value of this distance is proportional to *d*_*ISI*_ and is inferior to 1 when the considered trains match on average better than their shuffled versions. For all analyses, the timing-sensitive distance was computed using 1000 shuffling for each train (K = 1000). As illustrated in Supplementary Figure S6c-d, this timing-sensitive distance is now independent of the sum of the number of spikes in the two trains and of their difference. Moreover, the two distances *d*_*count*_ and *d*_*timing*_ are also uncorrelated (Supplementary Figure S7).

#### Population-wide spike train distances

Population-wide spike train distances were obtained by averaging normalized spike train distances across units:

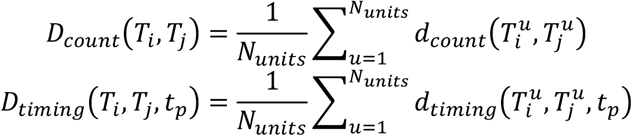

#### Temporal alignment of covert spike trains

When matching the trains occurring during covert speech to those occurring during the overt ‘repeat’ phase, the onset of the covert epoch was fixed to 600 ms after the appearance of the cross visual cue, which corresponded to the rounded average duration between the appearance of the visual cue and the speech onset in the repeat phase (true average duration of 596 ms when considering the first 31 trials).

#### Spike train classification of overt sentences using population-wide spike train distances

The population spike trains occurring during overt speech were classified with respect to 27 classes, corresponding to the uttered sentences. The classification used the (timing-based or count-based) population-wide spike train distance to perform basic pattern matching in a leave-one-out fashion. In each fold, the population spike train of one of the 62 utterances was isolated. The classification “model” consisted in the 61 remaining population spike trains labeled with the 27 classes. The population-wide distance was computed between the population spike train to be classified and all population spike trains of the “model”. The population spike train was attributed the label corresponding to the lowest population-wide distance.

#### Spike train classification of covert sentences using population-wide spike train distances

The population spike trains occurring during covert speech were classified with respect to 27 classes, corresponding to the different imagined sentences. The classification used the (timing-based or count-based) population-wide spike train distance to perform basic pattern matching with respect to the population spike trains occurring during the overt repetition. Each of the population spike trains of the 31 covert epochs was matched to the 31 overt repetition trains labeled with the 27 classes. The population-wide distance was computed between the population spike train to be classified and all population spike trains of the “model”. The population spike train was attributed the label corresponding to the lowest population-wide distance.

#### Unit selection for spike train classifications

For both overt and covert classification, only the units having more than 1 spike in at least 2 of the considered epochs were kept: One unit was excluded for overt classification and none for covert classification.

## SUPPLEMENTAL INFORMATION TITLES AND LEGENDS

**Supplementary Table 1.** List of the sentences and vowel sequences from the BY2014 corpus presented to the participant in the 31 considered trials.

| Trial | Sentence |
| --- | --- |
| 1 | a. i. ou. |
| 2 | u. é. è. |
| 3 | eu. o. an. |
| 4 | on. in. |
| 5 | Il y en a un. |
| 6 | Eux sont en guerre. |
| 7 | Nous sommes en mai. |
| 8 | C'est tout un monde. |
| 9 | Les temps ont changé. |
| 10 | Ils fuient le pays. |
| 11 | Au fond peu importe. |
| 12 | Saut en hauteur femmes. |
| 13 | Ce n'est pas un hasard. |
| 14 | La laïcité en actes. |
| 15 | La vague s'est étendue. |
| 16 | Une jeep a pris feu. |
| 17 | Voilà où nous en sommes. |
| 18 | Le lieu attend son heure. |
| 19 | Ou plutôt à ses manières. |
| 20 | Le quinze de France. |
| 21 | Le jeu est simple. |
| 22 | Son goût est sûr. |
| 23 | Cette guerre fut chère. |
| 24 | Il en paye le prix. |
| 25 | a. i. ou. |
| 26 | u. é. è. |
| 27 | eu. o. an. |
| 28 | on. in. |
| 29 | Il faut donc innover. |
| 30 | C'était nouveau et intéressant. |
| 31 | Il y aura une loi. |

**Supplementary Figure S1.**
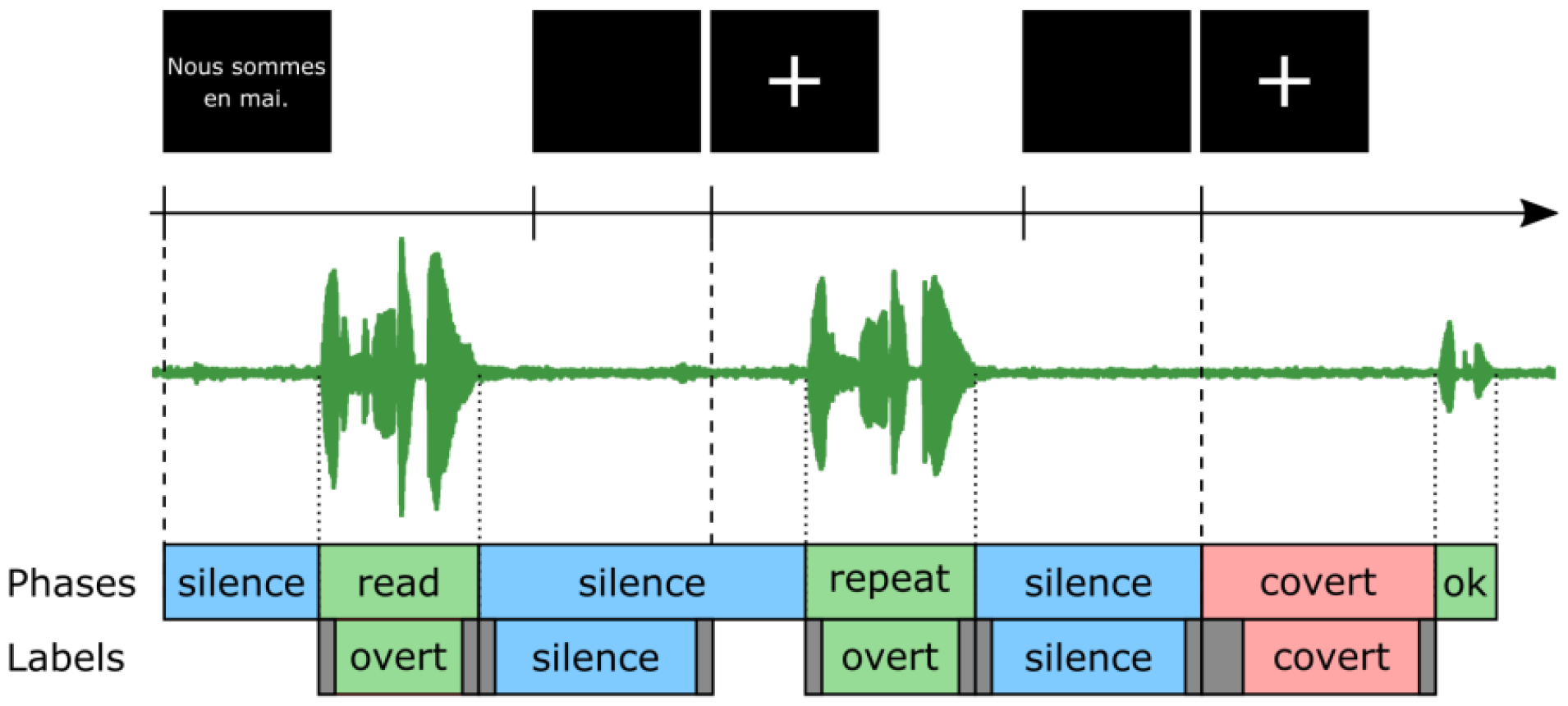
Course of a trial. The black rectangles represent the visual cues that were presented to the participant. The timeline indicates the time of appearance of the visual cues (see Methods). The green curve represents the audio signal. The first lines of colored boxes indicate how the trials were divided into phases corresponding to the different cognitive tasks performed by the participant. The second line indicate the labeled intervals that were considered for the classification of neuronal activity. On that line, the grey boxes show the margins that were excluded from each interval. All margins were 300-ms long, except for the covert-labeled epochs from which 600 ms were excluded at the beginning.

**Supplementary Figure S2.**
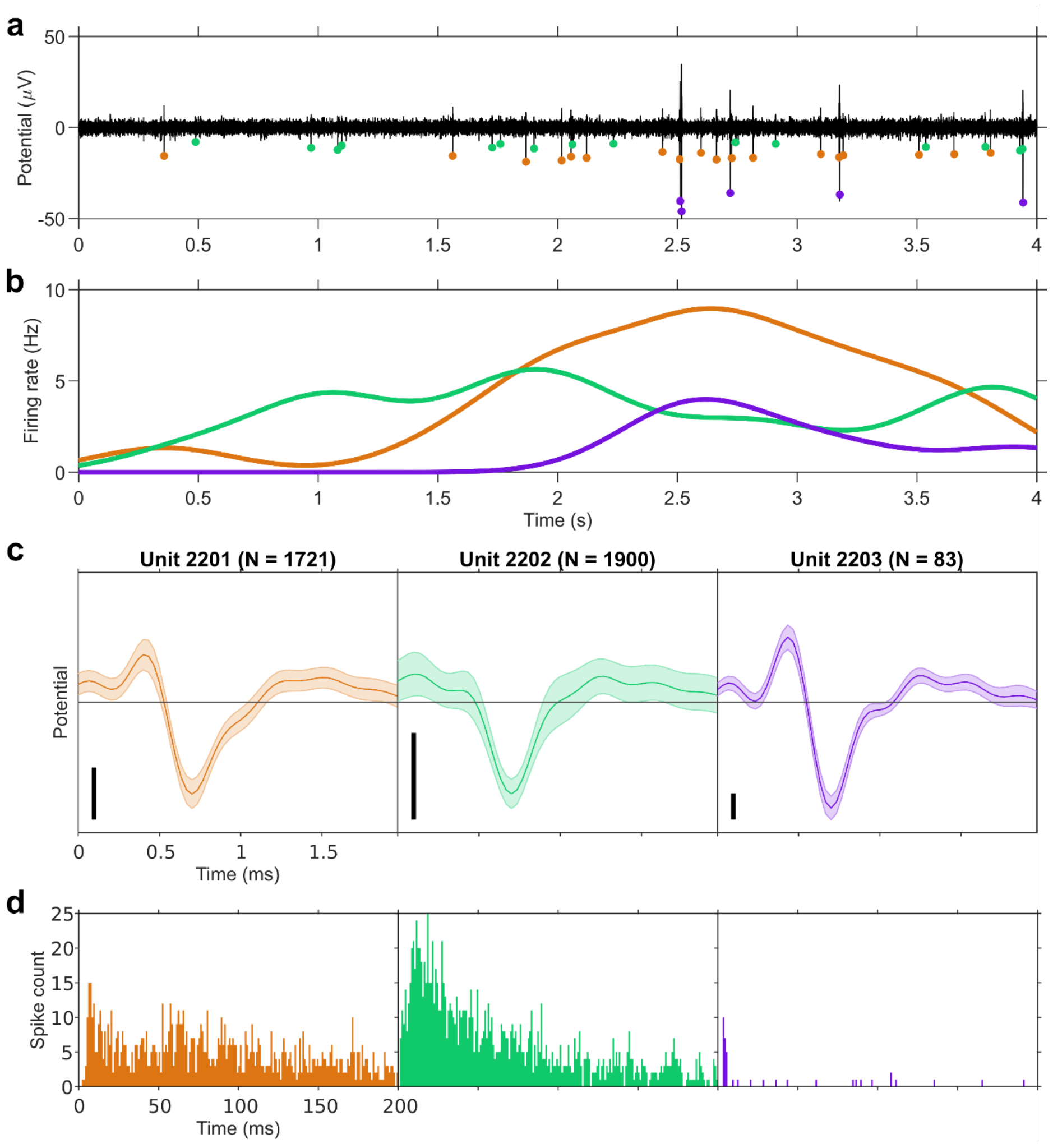
Example of spike sorting results and smoothed firing rates. **a. Four-second** extract of filtered signal from channel 22. The colored dots indicate detected spikes corresponding to the 3 units that were sorted for this channel, further shown in panel c. **b**. Smoothed firing rates of these 3 units along the extract displayed on panel a. **c**. Average waveforms of the 3 units sorted on channel 22. The scale bars represent 10 μV. The shaded area represents the standard deviation. N indicates the number of spikes detected for each unit. **d**. Distributions of the inter-spike intervals for the same 3 units as in panel a.

**Supplementary Figure S3.**
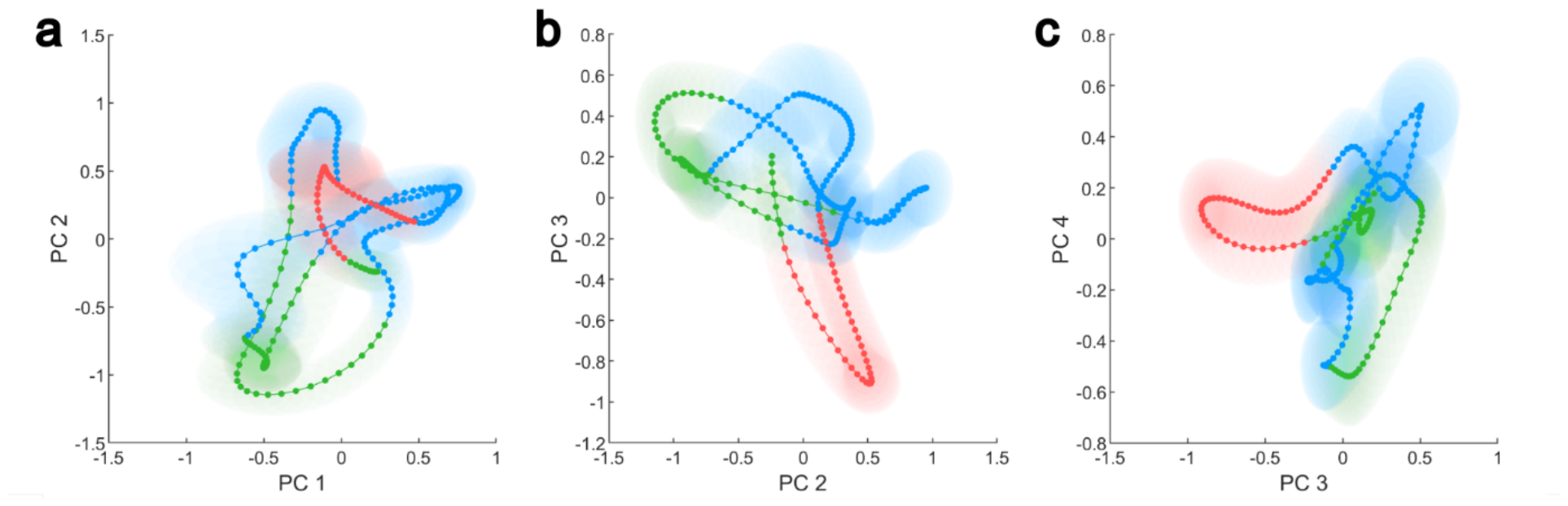
Trial-averaged projection of the firing rates of condition-modulated units using PCA. The firing rates of the 9 modulated units were projected in 2-dimensional spaces using PCA. The projected trajectories were averaged while preserving the different phases of the trial (see Supplementary Figure S1). The shaded areas are semi-transparent disks whose radius is equal to the standard error of the mean (SEM). SEM was chosen instead of standard deviation in order to improve readability. **a** Projection on PCs 1 and 2. **b** Projection on PCs 2 and 3. **c** Projection on PCs 3 and 4. Colors as in Figure 2b: green=overt, blue=silence, red=covert.

**Supplementary Figure S4.**
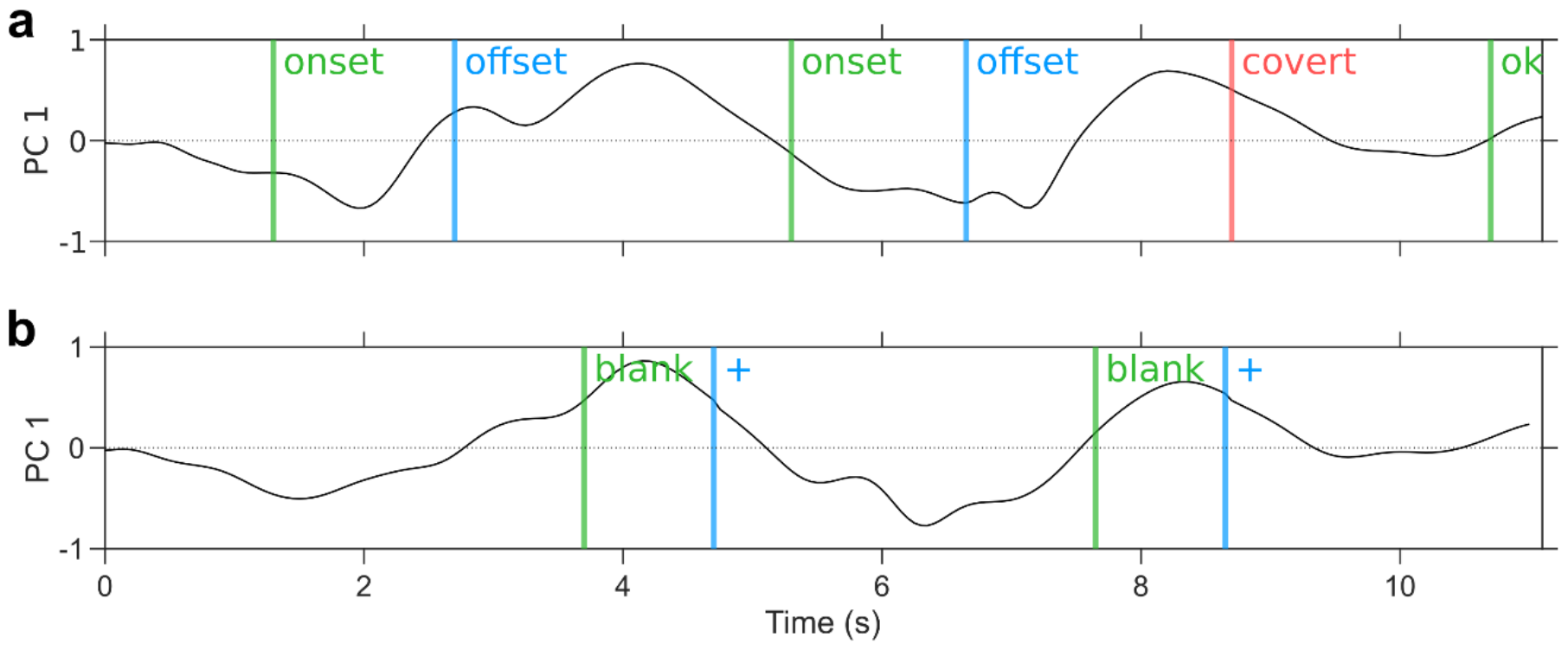
Trial-averaged first principal component of the firing rates of condition-modulated units with different alignments. **a** Average of first principal component when aligned on trial phases. **b** Average of first principal component when aligned on visual cues. See also Supplementary Figure 1.

**Supplementary Figure S5.**
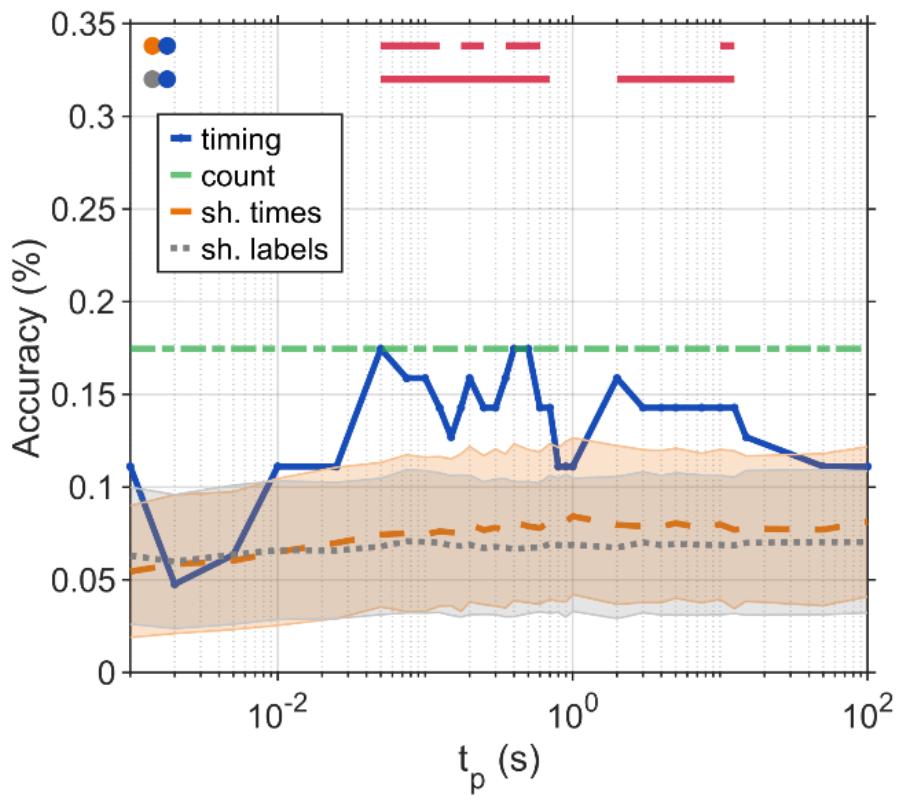
Classification of overtly produced sentences from ensemble spiking activity as in Figure 4a but when considering surrogate time-shifted overt read and repeat windows. Compared to Figure 4a, here the read and repeat windows are shifted back in time so that for each trial, the surrogate read window ends at the beginning of the trial, when the sentence is displayed. This procedure ensures that the subject had not yet seen on the screen the sentence during the surrogate read window.

**Supplementary Figure S6.**
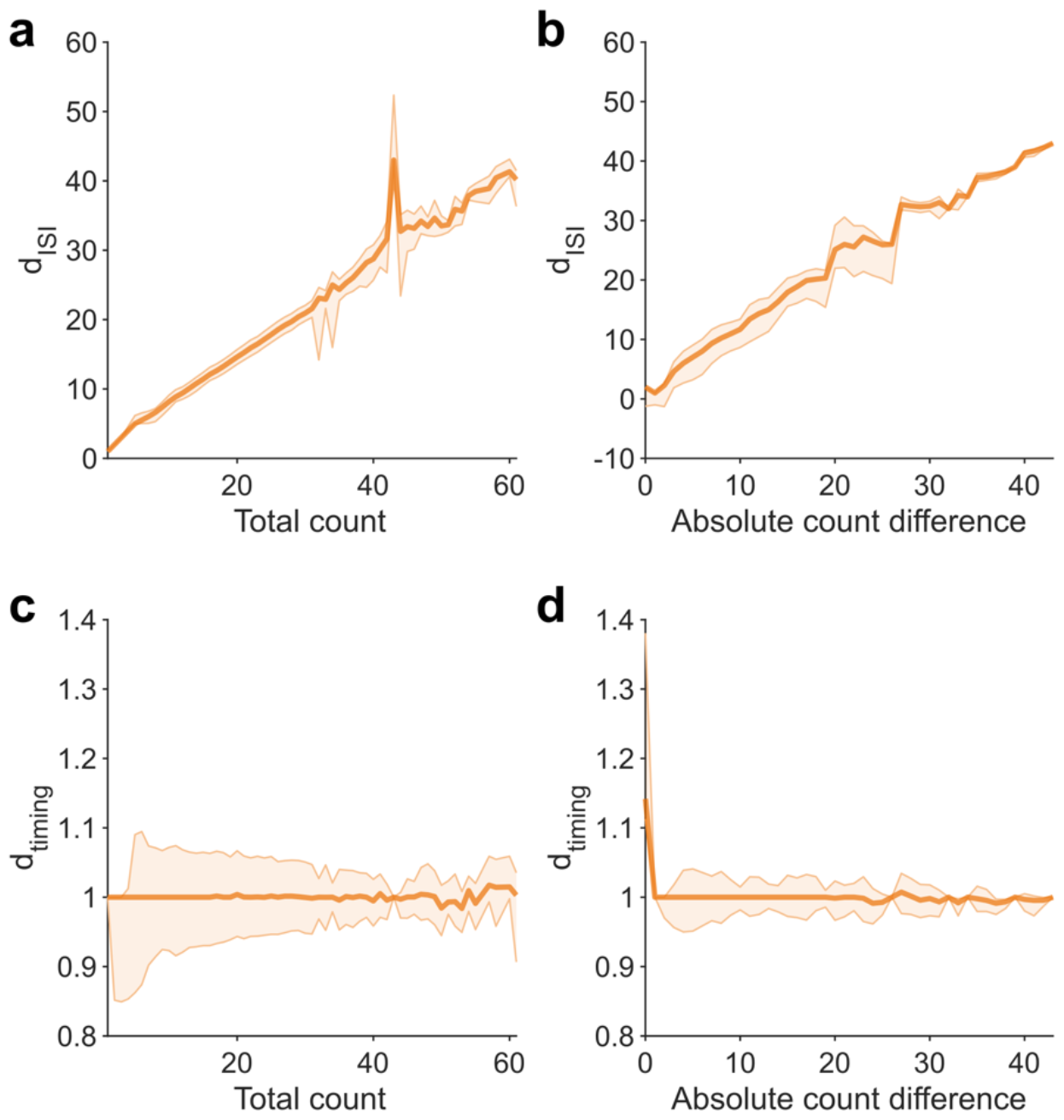
Effect of normalization of the original VP distance on its correlation with spike counts. **a-b**. The original VP distance strongly correlates with spike counts and difference in spike counts between both trains. **c-d**. The modified VP distance d_timing_ no longer correlates with spike counts and difference in spike counts. The measures were obtained using 10 temporally shuffled versions of the spike trains extracted for the overt classification process. For each temporal shuffling, the distances between all pairs of spike trains were computed. For each panel the solid line indicates the median and the shaded area the 25^th^ and 75^th^ percentiles across all shufflings and pairs of spike trains.

**Supplementary Figure S7.**
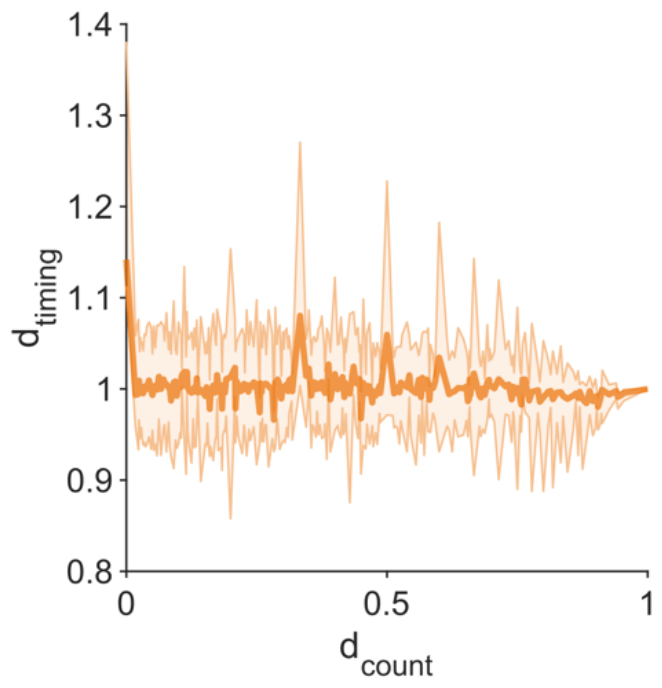
Normalized timing-based distance as a function of the count-based distance, showing that both measures are uncorrelated (see Methods). The measures were obtained using 10 temporally shuffled versions of the spike trains extracted for the overt classification process. For each temporal shuffling, the distances between all pairs of spike trains were computed. The solid line indicates the median and the shaded area the 25^th^ and 75^th^ percentiles across all shufflings and pairs of spike trains.

